# Tracking tendon fibers to their insertion – a 3D analysis of the Achilles tendon enthesis in mice

**DOI:** 10.1101/2020.02.25.964759

**Authors:** Julian Sartori, Heiko Stark

## Abstract

Tendon insertions to bone are heavily loaded transitions between soft and hard tissues. The fiber courses in the tendon have profound effects on the distribution of stress along and across the insertion. We tracked fibers of the Achilles tendon in mice in micro-computed tomographies and extracted virtual transversal sections. The fiber tracks and shapes were analyzed from a position in the free tendon to the insertion with regard to their mechanical consequences. The fiber number was found to stay about constant along the tendon. But the fiber cross-sectional areas decrease towards the insertion. The fibers mainly interact due to tendon twist, while branching only creates small branching clusters with low levels of divergence along the tendon. The highest fiber curvatures were found within the unmineralized entheseal fibrocartilage. The fibers inserting at a protrusion of the insertion area form a distinct portion within the tendon. Tendon twist is expected to contribute to a homogeneous distribution of stress among the fibers. According to the low cross-sectional areas and the high fiber curvatures, tensile and compressive stress are expected to peak at the insertion. These findings raise the question whether the insertion is reinforced in terms of fiber strength or by other load-bearing components besides the fibers.

## 1 Introduction

Entheses are regions within the locomotor system where tensile forces are transmitted between dense regular connective tissues, like tendons or ligaments, and bone. The stiffness of bone, approximated by Young’s modulus [1], is one to two orders of magnitude higher than that of tendon [2]. Any mismatch in deformation behavior can cause additional stress when the interface is subjected to tension [3]. The research question behind this study is whether and how the enthesis achieves a homogeneous distribution of stress along and across the insertion to bone, in spite of changing angles in the corresponding joints.

### 1.1 The Achilles tendon in mice

The Achilles tendon in mice connects the three muscle bellies of the *M. triceps surae* to an insertion area on the posterior side of the *Calcaneus*. Posterior crural fascia and *Tuber calcanei* deflect the tendon course to an s-shape. While the cross-sectional shape of the tendon is elliptical at mid-length, it gets flattened adjacent to the *Tuber calcanei* [4].

The fiber courses within the Achilles tendon are twisted in many species [5,6]. From an ultrastructural perspective fibers are fibril bundles. They can, therefore, branch into smaller bundles or fuse into larger ones [7]. Fibrils can be crimped with varying crimp periods and angles [8,9]. The crimps are usually aligned within fibers [10]. The fibers dissociate at the interface with bone. But in the porcine anterior cruciate ligament, the fibrils go on [11]. On the fibrillar level tendon and bone could be continuous.

The insertion of the Achilles tendon to the *Calcaneus* is a fibrocartilaginous enthesis [12,13]. The transition between tendon and bone does not occur abruptly. Instead, the collagen fibers pass through unmineralized and mineralized fibrocartilage before reaching the bone. Furthermore, the region of the tendon that is in contact with the *Tuber calcanei* is a sesamoid fibrocartilage [13]. The fibrils in tendon tissue mainly consist of type I and III collagen [10], in the fibrocartilages a higher share of collagen type II is found [14–17]. Importantly, the glycosaminoglycan content of fibrocartilage is much higher than that of tendon tissue [16,18]. Glycosaminoglycans bind high amounts of water. The matrix between the fibers, therefore, is more compression-resistant. The transition from unmineralized fibrocartilage to mineralized fibrocartilage is characterized by a gradual increase in mineral content within and between the fibers [19].

### 1.2 Questions on the fibrous structure

This study of the Achilles tendon enthesis in mice was carried out to understand the mechanics of the enthesis. The preceding description shows that tendon fibers are diverse in properties. Many of these properties have an effect on mechanical behavior. Therefore, fibers of the Achilles tendon were tracked from a position in the free tendon to the insertion in volume images from micro-computed tomography (µCT). Fiber tracks and slices from the µCT-data were analyzed to elucidate the following aspects of the fibrous structure.

1. The sum of the fiber cross-sectional areas (CSA) was found to decrease towards the insertion in a previous study [4]. The finding is counter-intuitive because it implies increased tensile stress near the interface. We do not know whether this is due to a decrease in fiber number or in CSA per fiber. We hypothesize that fiber numbers stay about constant along the tendon and that the cross-sections of fibers are compressed against the *Tuber calcanei* in the distal tendon. The CSA of single fibers should hence decrease towards the insertion.
2. Fiber branching and tendon twist could contribute to a homogeneous distribution of stress among the fibers [6] in spite of changing angles in the joints. Branching also could increase the shear stiffness of the tendon and prevent fibers from sliding relative to one another. Therefore, the fiber tracks were examined with regard to branching points and clusters formed by branching fibers. To measure twist, the angular deviation of fiber tracks from the tendon course was measured at regular intervals along the tendon.
3. Fibrocartilages are considered an adaptation to compressive loads [15,20]. The pressure exerted by a fiber on a neighboring rigid structure can be approximated by the tensile force, the width of the fiber and its curvature around that structure [21]. Therefore, we quantified the fiber curvature to examine whether regions with high curvatures correspond to the fibrocartilages. Furthermore, we approximated the contact pressure exerted by the fibers.
4. The insertion area on the *Calcaneus* is not completely smooth in mice. Parts of the mineralized fibrocartilage form a protrusion. The deep fibers of the Achilles tendon insert in the prominent surface of the protrusion. The superficial fibers cover this part of the enthesis and insert more distally drawing an acute angle with the hard tissue surface. A possible cause for the formation of the protrusion could be a different loading regime. Therefore, the fibers of the protrusion were tracked to find out whether they correspond to a distinct portion of the more proximal tendon or spread over the complete tendon area.

### 1.3 Fiber tracking techniques

For fiber tracking, it is necessary to identify the fibers in the surrounding tissue. A number of methods exist for this purpose, which can be divided into destructive and non-destructive methods. Destructive methods include, for example, layerwise preparation and serial sections [22–25]. In contrast, the non-destructive methods can be classified according to their modalities, i. e. X-ray tomography, magnetic resonance tomography and 3D ultrasound [26–29]. For most methods, the data acquisition is followed by digital image processing, i.e. stereology, edge detection, template-based fiber tracking, Fourier analysis, and texture analysis. After determining local directional properties, fiber tracks are reconstructed starting from so-called seed points. An exception is the diffusion tensor imaging technique, since here the fiber orientation required for fiber tracking even is recorded by the imaging modality [30,31].

With these methods, muscle fibers have been analyzed using texture-based [26,28] and template-based fiber tracking [32]. Collagen fibers were tracked by Disney et al. [27] in the *Annulus fibrosus* of bovine intervertebral discs. They scanned their specimens with propagation-based phase-contrast X-ray tomography at high resolutions (0.81 µm voxel size), carried out template-based fiber tracking and examined the distribution of fiber orientations. This is the first analysis of the three-dimensional fiber tracks in macroscopic tendon samples. Earlier publications have reported the segmentation of tendon fascicles in macroscopic tendon samples [33,34]. Fibers [7] and fibrils [35–37] have been analyzed in differently sized subvolumes of tendons.

## 2 Material and methods

### 2.1 Animals, sample preparation and imaging

This study is a detailed analysis of volume images of three hind limbs (two left hind limbs and one right hind limb), each from another mouse individual (*Mus musculus*, strain C57BL/6J, all-female and aged 7.5 months). We chose to examine the Achilles tendon in mice because in small animals the ratio of fiber size to spatial resolution in a µCT is maximal. At the same time, the literature about the Achilles tendon enthesis in mice is comprehensive enough to provide sufficient context. The euthanasia and dissection of the three mouse individuals, as well as the treatment and imaging of the hind limbs, are described in detail in a previous article [4]. Briefly, the hind limbs were fixed in formalin, cell- macerated [38], demineralized and cropped to a central portion keeping the *M. triceps surae* intact from origin to insertion. The resulting specimens were dried over-critically and scanned in a µCT setup (ZEISS Xradia 520 Versa by Carl Zeiss AG, Oberkochen, Germany) with low tube voltages and propagation-based phase contrast. The parameters were set as follows: 30 kV source voltage, 4× optical magnification, 16 mm source–sample distance, 25 mm sample–detector distance, 70 s exposure time and 3,201 angular steps. All procedures involving animals conformed to national and international ethical standards.

### 2.2 Fiber tracking

The isometric voxel size of the volume images is 1.32 µm. The gray values are 16-bit encoded. The volume images were segmented manually in a 3D image analysis software (Amira Software 2019.1 by Thermo Fisher Scientific Inc., Waltham, Massachusetts, USA). Tendon, bone, the insertion area and the protrusion of the insertion area were marked as segments.

The fibers were tracked within the tendon segments with a template-based fiber tracking procedure (Amira Xtracing Extension 2019.1) involving the following two steps. (1) At each voxel position, a cylinder template with a given geometry is rotated with a defined angular step size. For each rotational step, the correlation with the volume data is measured. The orientation vector with the maximal correlation is saved in an orientation field, the corresponding correlation value in a correlation field. The cylinder correlation analysis was calculated with the following parameters: cylinder length 30 µm, angular sampling 5°, mask radius 5 µm, cylinder radius 4 µm, bright fibers on dark background. (2) The fiber tracking module starts fiber tracks at positions where the correlation is above a threshold (“maximum seed correlation”). It searches for candidate points within a search cone defined by the orientation vector. The fiber tracking stops when the correlation value of all candidate points weighted by the distance from a straight line is below another threshold (“minimum correlation quality”). The input parameters were selected as follows: Minimum seed correlation 170, minimum continuation quality 100, direction coefficient 0.3, minimum distance 8 µm, minimum length 30 µm, search cone length 30 µm, search cone opening angle 37°, minimum step size 5 % of search cone length. The result is one set of multi-segmented lines (fiber tracks) per specimen. These sets were pruned to the tendon segment. Fiber tracks with less than two line segments were removed.

### 2.3 Spatial reference system

A mid-tendon line has been calculated previously for each specimen [4]. It is as well a multi-segmented line. The lengths along each mid-tendon line beginning at the insertion were normalized with *Calcaneus* height to obtain normalized tendon length (NTL) positions that are comparable between the specimens. The *Calcaneus* heights of the specimens were 949 µm, 957 µm and 999 µm. The extraction of transversal tendon sections from the datasets was implemented as an Amira script. The sections were extracted at positions along the mid-tendon line with NTL intervals of 0.05 *Calcaneus* heights between neighboring sections. The normal vector of the section plane is defined by the tangential vector of the mid-tendon line at this position. The first transversal section is extracted at the intersection of the mid-tendon line and the tidemark of the Achilles tendon insertion at the *Calcaneus* (NTL = 0.00). The examined length of the tendon ends at a NTL of 1.25 to 1.45 depending on the specimen. In spite of this definition of NTL, the terms ‘‘proximal’’ and ‘‘distal’’ are strictly used with reference to the trunk.

### 2.4 Analysis of fiber tracks and transversal sections

For the following analyses, the images were processed with Amira, ImageJ (ImageJ 1.52p by National Institutes of Health, Bethesda, Maryland, USA) and the custom software ImageXd (version 2.9.25) and Cloud2 (version 14.11.29). ImageXd and Cloud2 are programmed by one of the authors (HS, https://starkrats.de). Analyses were carried out with Amira, Cloud2, and ImageJ. Volume datasets and sets of fiber tracks were rendered with Amira.

#### 2.4.1 Fiber counts and measures

Two approaches are compared to measure the number of fibers. (1) The number of fibers was determined from the fiber tracks. Therefore, the number of tracks intersecting any of the defined transversal section planes was determined in each specimen. (2) Sections were extracted from the original volume images and binarized with a threshold (gray value of 18,500). The resulting masks were further subdivided with binary watershed segmentation and counted with the particle analysis tool within ImageJ.

Four further measures were extracted from the particle analyses: the areas, maximal and minimal Feret’s diameters of the fiber masks in the sections and the nearest neighbor distances of their centroids (nearest neighbor distance plugin by Yuxiong Mao). The distribution of each of these measures is reduced to the median, the first and the third quartile for each section. These analyses were carried out for all three specimens (Fig. S2A, B). For the sake of clarity, results for one specimen are presented.

#### 2.4.2 Endpoints of fiber tracks and branching

The proximal and distal endpoints of the fiber tracks were extracted and counted within lengthwise tendon segments to analyze the lengthwise distribution of fiber terminations. The lengthwise tendon segments were generated by a Cloud2 script subdividing each set of endpoints at the transversal sections. A random sample of 42 distal endpoints from one specimen was evaluated visually. Fiber tracks and volume data were cropped to a sphere around each endpoint, visualized together (Fig. S1A) and inspected. Depending on the arrangement of the fibers and the position of the sphere in the volume data, the endpoint was assigned to one of the following classes: (1) fiber inserts to hard tissue, (2) fiber branches into another fiber, (3) fiber leaves the tendon, (4) fiber factually ends within the tendon and (5) fiber track corresponds to tracking artifact.

The branching pattern was approximated in one specimen by merging fiber tracks and endpoints whose distance was below a threshold distance of 8 µm or equals it. This approximation is supported by the minimal distance criterion of the tracking algorithm and the visual evaluation of endpoints. The fiber tracks were refined to a step size of 1 µm on the basis of polynomial fits for this calculation. The resulting clusters of branching fiber tracks were further evaluated. The number of fiber tracks per cluster was counted and a random sample of clusters was exported and compared to the volume data. For this comparison, the single fibers in the volume data were automatically segmented. The fiber tracks were used as markers for this segmentation.

### 2.4.3 Orientation and curvature of fiber tracks

The angular deviations of the fiber tracks from the mid-tendon line are measured in lengthwise tendon segments. To this end, the fiber tracks were subdivided at the transversal sections in the way described for the endpoints. The angular deviation is defined as the angle between the fiber track and the mid-tendon line in a lengthwise tendon segment. Each fiber track and the mid-tendon line were reduced to a vector connecting the first and last points within the segment for this measurement. The distribution of the deviation angles is reduced to the median, the first and the third quartile for each segment. These analyses were carried out for all three specimens (Fig. S2C). For the sake of clarity, results for one specimen are presented.

A polynomial (maximal degree 7) was fit to each fiber track in one specimen in Cloud2. Refined fiber tracks with a maximal angular deviation of 1° from the polynomial were generated to measure the local curvature radius. The radius values were smoothed by calculating the local average. They were saved in the additional parameter *t* at each point of each fiber track.

#### 2.4.4 Region of insertion and length reserves

Tracks of fibers inserting at the protrusion of the enthesis were labeled with a module within Amira (“Label spatial graph”) and their distribution was inspected at the proximal end of each set of fiber tracks. Tortuosity, the relation between curved length and chord length, was only measured in long fiber tracks reaching from the insertion to the proximal end of each dataset. It was measured in each track of these subsets by a module within Amira (“Spatial graph statistics”) and is given as a percentage indicating how much longer the curved length is relative to the chord length.

#### 2.4.5 Estimation of stress

Tensile stress in the fibers is calculated from the sum of the fiber cross-sectional areas at a NTL of 0.20 and 1.25 in one specimen and from the maximal isometric force of the plantar flexors (F = 3.3 N, value involves *M. plantaris*) [39]. For this calculation, we assume that the force is only transferred by the fibers and that stress is homogeneously distributed over CSA of the fibers. The tensile force transferred by a single fiber is calculated under the same assumptions from the fiber number and the maximal isometric force. The contact pressure exerted by a single fiber is approximated using the model of a strap pressing against a pulley [21]. The calculation is based on the assumption that bending and shear stiffness of the strap, as well as the friction between strap and pulley, are negligible. Contact pressure is calculated from the maximal Feret’s diameter, the fiber radius and the tensile force in the fiber.

## 3 Results

### 3.1 Fibrous structure and fiber tracks

Fibers can be visually identified in the volume images by their high gray values and unidirectional elongation along the tendon. The determined fiber tracks are located in the centers of such fibers. The fiber tracks agree among the three specimens with regard to their courses. They are roughly aligned with the tendon course but exhibit a low level of twist around the tendon center. Endotenon and a subdivision into fascicles were not found in any of the specimens.

### 3.2 Fiber numbers

The mean fiber number as determined by the particle analysis of binarized sections amounts to approximately a thousand fibers at NTLs between 0.20 and 1.25 (Fig. 1A–D). The fiber number steeply decreases from a NTL of 0.20 to 0.00. According to the volume renderings (Fig. S1D), fibers insert into hard tissue in the region corresponding to a NTL below 0.18.

**Fig. 1:**
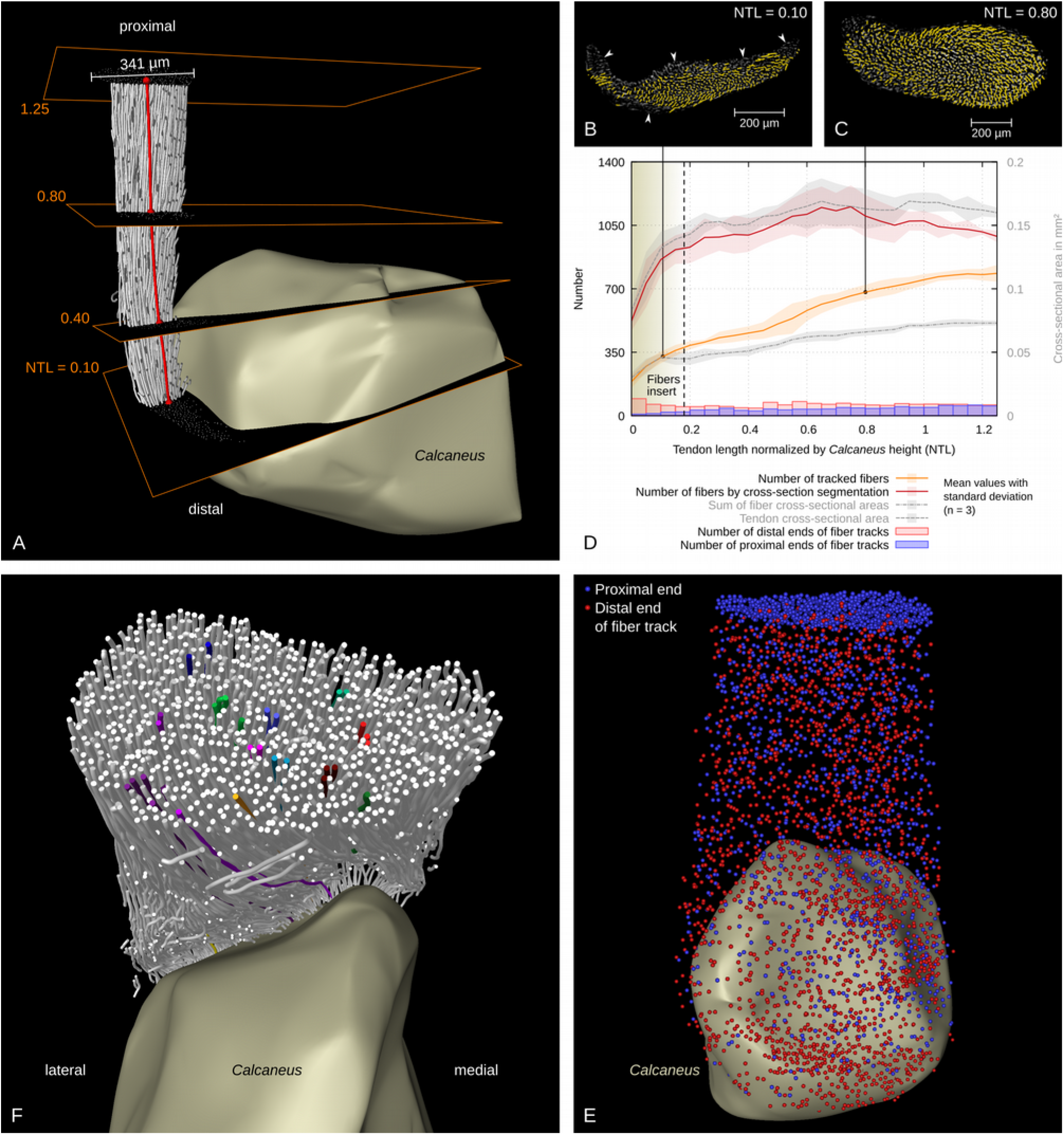
Fiber numbers and branching. (A) Rendering of the fiber tracks at the insertion of the right Achilles tendon in a mouse. Transversal sections defined by the mid-tendon line (red) as normal vector were extracted from the volume data at regular normalized tendon length (NTL) intervals along the mid-tendon line. The fiber tracks were compared to the volume data, i.e. in (B) a distal section close to the insertion, where no fiber tracks were found in some regions of the cross-section (arrowheads), and in (C) a proximal section. (D) Fiber numbers were determined by a particle analysis of such sections after segmentation (red curve) and by counting the fiber tracks crossing the sections (orange curve). A NTL (x-axis) of zero corresponds to the insertion of the mid-tendon line. At positions left of the vertical dashed line fibers insert to the Calcaneus. The tendon CSA and the sum of the fiber CSAs are plotted for comparison [4]. Furthermore, the number of proximal and distal endpoints of fiber tracks in each volume segment is plotted. (E) Rendering of proximal (blue) and distal endpoints of fiber tracks (red). (F) Rendering of several clusters of fiber tracks formed by branching.

The mean fiber numbers determined from the fiber tracks are much lower. They decrease from 785 (80 % of the values from the particle analysis) in the most proximal section to 388 shortly above the insertion zone (NTL = 0.2). The regions, where no fiber could be tracked, were larger in the distal than in the proximal tendon (Fig. 1B, C). In particular, fiber tracks were not identified in fibrocartilaginous regions where the density of transversal connections between the fibers is high (Fig. 3D, open arrows). The automatic segmentation of single fibers attributes large regions to a few fibers in the distal tendon (Fig. S1C).

### 3.3 Fiber sizes and distances

Fiber sizes and distances were measured in binarized transversal sections of the µCT-data (Fig. 2A–F). The medians of the minimal and maximal Feret’s diameters decrease from proximal to distal positions along the tendon and slightly increase again within the enthesis. The median of the maximal Feret’s diameter ranges between 10 µm and 13 µm. The ratio of the minimal to the maximal Feret’s diameter stays about constant over the examined length. It ranges between 0.59 and 0.63. The median CSA of the fibers decreases from 62 µm^2^ in the proximal tendon to 39 µm^2^ shortly above the insertion (NTL = 0.25) and slightly increases within the insertion.

**Fig. 2:**
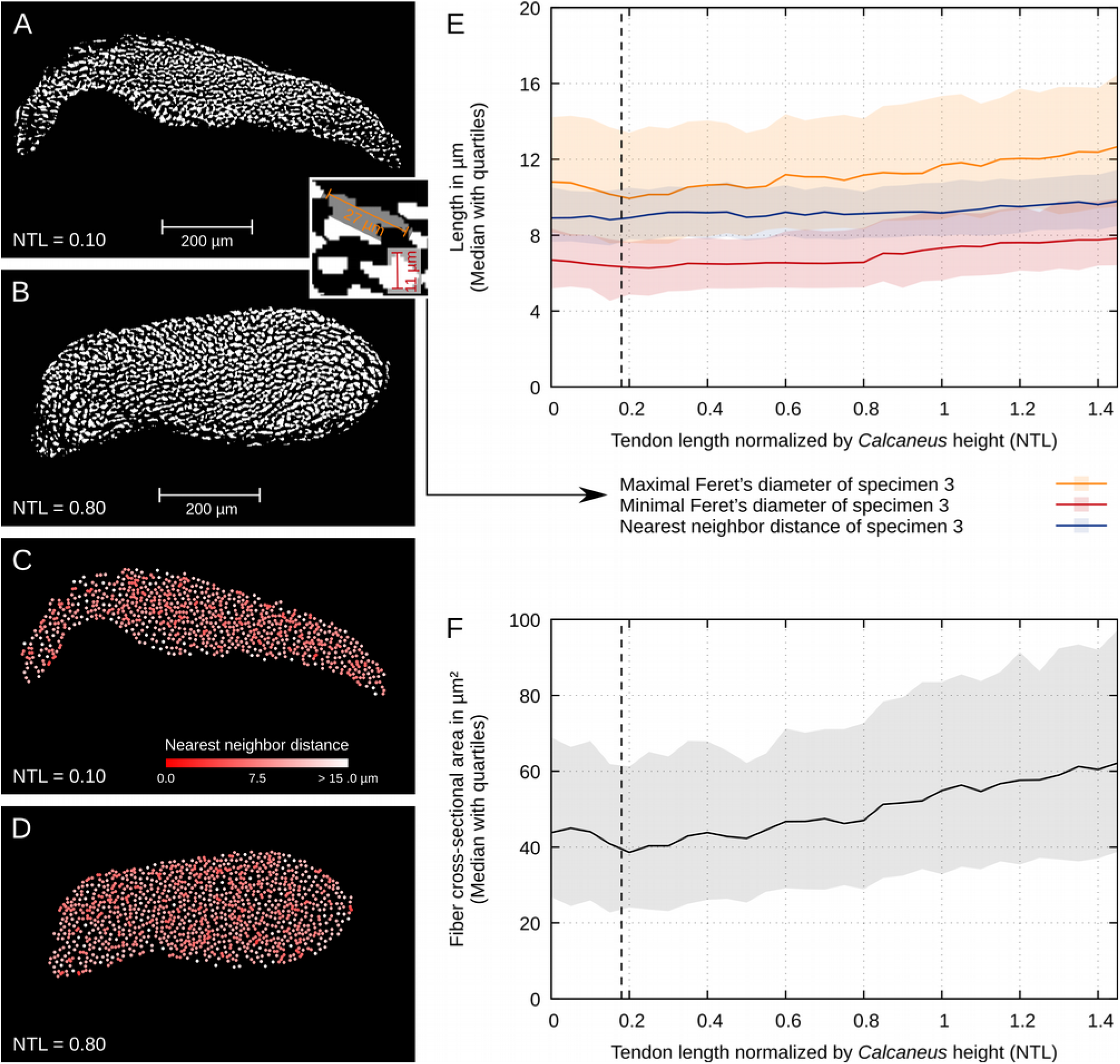
Fiber cross-sectional shape, distance, and size. (A) Binarized and segmented sections through the distal tendon at a NTL of 0.10 and (B) the more proximal tendon at a NTL of 0.80. The inset illustrates how maximal and minimal Feret’s diameter was measured. (C) The distribution of the nearest neighbor distances between the centers of the identified fiber sections is homogeneous within sections through the distal tendon and (D) the more proximal tendon. (E) Both median Feret’s diameters increase with distance from the insertion, while the the median nearest neighbor distance stays about constant. (F) Accordingly, the median CSA per fiber increases with distance from the insertion.

The nearest neighbor distances are homogeneously distributed within any of the cross-sections. Their section-wise median only decreases slightly from proximal to distal. It ranges between 8.8 µm and 9.8 µm. Together with the decreasing fiber size, this implies that the gaps between the fibers become larger from proximal to distal because the nearest neighbor distances were measured between the centroids of fiber masks.

### 3.4 Fiber branching

The visual inspection of distal endpoints of fiber tracks (n = 42) shows that none corresponds to a factual fiber end. The tracking procedure cannot identify fiber branching. Instead, one of the branches will be rendered as a fiber track that ends at the branching point. 52 % of the inspected endpoints correspond to such branching points (Fig. S1B). In 12 % of the endpoints, the corresponding fiber passes over into hard tissue. In 5 % the fiber leaves the tendon segment at another boundary. 29 % were related to tracking artifacts. We noted a position where fiber kinking had occurred in one specimen. This led to interruptions in the fiber tracks.

Endpoints of fiber tracks only occur at higher densities at the proximal end of the dataset, near the insertion and at the described kinking artifact. Otherwise, they are distributed homogeneously over the tendon volume (Fig. 1D, E).

The dataset that was used for reconstruction of branching topology comprised 1,982 fiber tracks. They cluster in 1,159 entities with the following distribution of cluster sizes: 71 % of the resulting entities correspond to single fibers that do not branch, 13 % correspond to branching clusters of two fibers, 7 % comprises three fibers, 9 % are distributed among clusters with four to thirteen fibers. Furthermore, there is one cluster with 17 and one with 24 fibers. The fiber tracks of most of the clusters do exhibit low levels of divergence and only extend over small shares of the tendon CSA (Fig. 1F).

### 3.5 Helical fiber courses

Renderings of the fiber tracks (Fig. 3D) from a right hind limb show that most fibers follow a left-handed helix. In the left hind limbs right handed helices where found. Some fibers in the tendon center do not undergo a helical twist but follow a rather straight course.

**Fig. 3:**
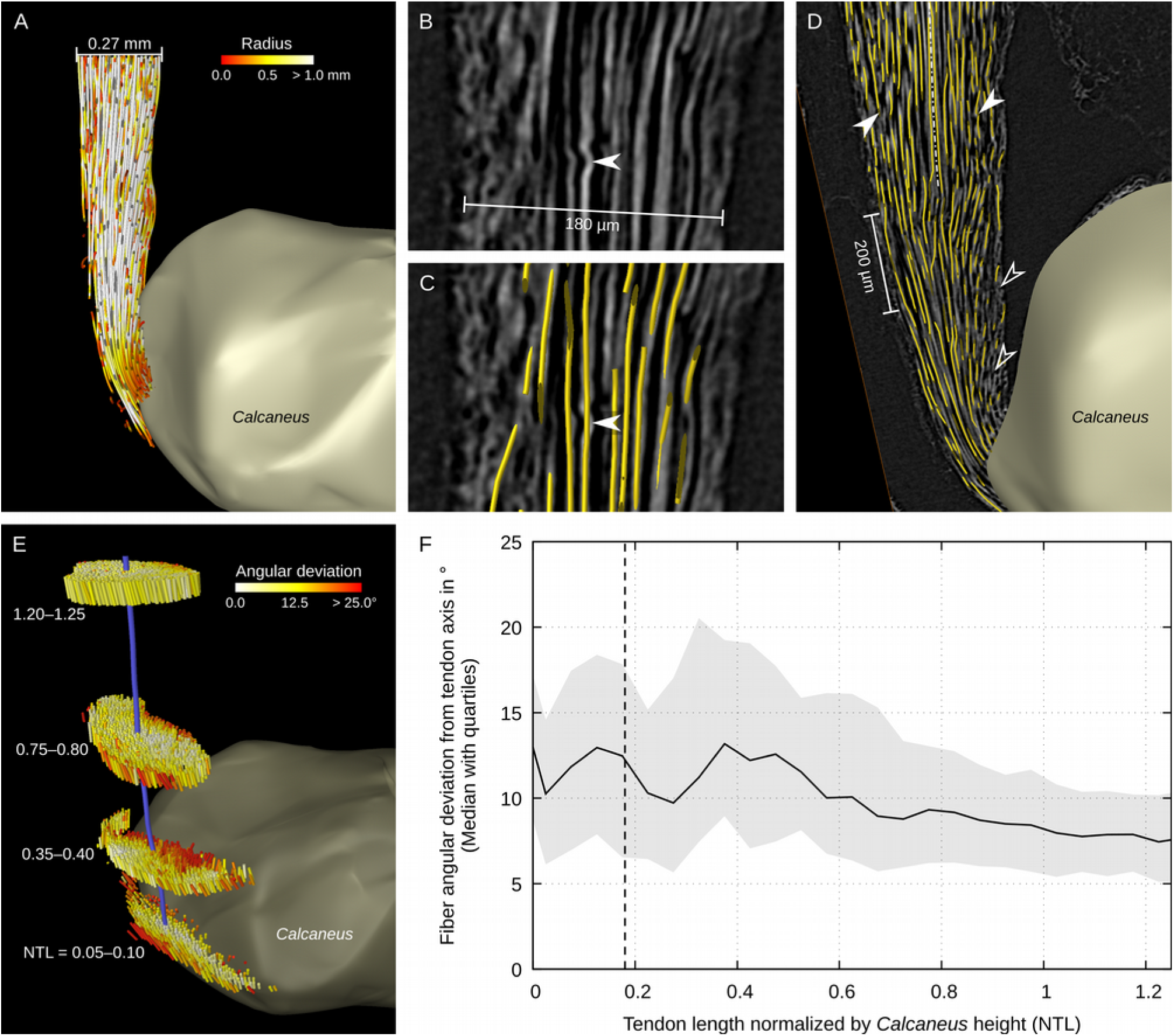
Fiber curvature, twist, and orientation. (A) Rendering of the fiber tracks cut at a sagittal slice. The colors indicate the local radius. Fiber curvature is high near the insertion. (B) Undulations of fibers are not reflected in (C) the fiber tracks. Length measurements, therefore, do not include fiber crimp. Crimp period of the shown undulation is 22 µm, crimp angle is at least 19°. (D) Rendering of fiber tracks and of a sagittal slice through the volume data. While central fibers are exactly in the cutting plane, the ends of fiber tracks (filled arrowheads) indicate a left-handed helical course. In fibrocartilaginous regions with transversal connections between the fibers (open arrowheads), fewer fiber tracks were identified. (E) Rendering of the fiber vectors in lengthwise segments of the tendon. The colors indicate the angular deviation from the mid-tendon line (violet). The angular deviations are higher at the margins of the cross-sections. (F) The median angular deviation of fiber vectors from the mid-tendon line is plotted over the NTL. The median angular deviations near the insertion are higher and the distributions broader than in the proximal tendon.

Orientations were examined in the lengthwise tendon segments (Fig. 3E, F). The median angular deviation of the fiber tracks from the tendon axis is 7.5° in the proximal tendon. Distally, fibers become less aligned with the tendon axis. The median angular deviation reaches a local maximum of 13° above the insertion and exhibits equally high values within the insertion. Likewise, the variation of angular deviation is higher in the distal than in the proximal tendon. Within the cross-sections, the highest angular deviations are found at the anterior and posterior margins, while the fibers in the center of the tendon are more aligned with the tendon axis.

### 3.6 Distribution of fiber curvature

The fibers in the proximal tendon, in the tendon center, and at the posterior margin follow straight courses. High local curvatures with radius values below 0.3 mm are found in the enthesis and in the medial and lateral margin of the distal tendon (Fig. 3A). In some regions, local waviness of fibers can be seen in the volume data. The corresponding fiber tracks do not follow the undulations but their central axis (Fig. 3B, C).

### 3.7 Fibers of the protrusion

Long fibers are tracked from the regions of the insertion to the proximal end of the dataset (Fig. 4A, B). The fibers inserting at the protrusion correspond to just an anterolateral share of the proximal tendon cross-section. The fibers that fill the posterior part and the medial margin in the proximal tendon mainly insert at acute angles in a crescent area surrounding the protrusion distally. The tortuosity of the long fiber tracks ranges between 0 % and 15 %. The median tortuosity amounts to 1 %. The fibers of the protrusion tend to exhibit lower tortuosities than the surrounding fibers (Fig. 4C).

**Fig. 4:**
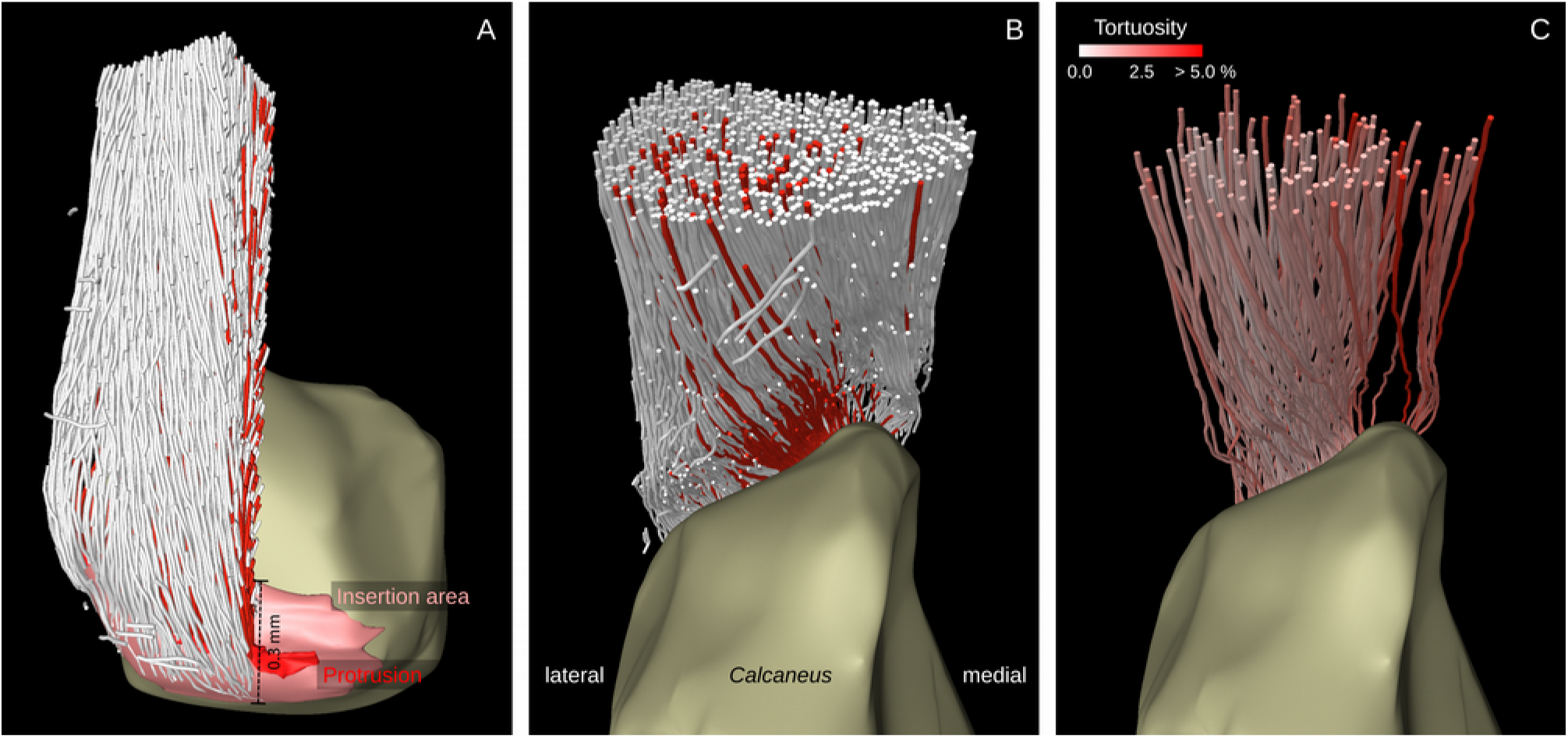
The fiber tracks ending at the protrusion (red surface) were (A) marked near the insertion area and inspected (B) from a proximal perspective. The fiber tracks are only distributed within an anterolateral share of the proximal tendon cross-section. (C) Tortuosity, the ratio of curved length and chord length, is measured in fiber tracks that reach from the insertion to the proximal end of the dataset. Fiber tracks from the protrusion tend to be more taut, while the fiber tracks in the posterior and medial margin of the tendon have higher length reserves.

### 3.8 Tensile and compressive stress

The estimated tensile stress amounts to 46 MPa in a proximal cross-section (NTL = 1.25) and to 80 MPa in a distal cross-section (NTL = 0.20). The force transferred by a single fiber is approximated to 3.2 mN in a proximal cross-section and to 3.9 mN in a distal cross-section. Curved fibers in the enthesis (NTL = 0.05) are estimated to exert a contact pressure of about 1.2 MPa.

## 4 Discussion

### 4.1 The fibrous structure is not reinforced at the enthesis

For measuring the fiber numbers a segmentation of sections is preferred over measurements from the fiber tracks because several factors complicate the identification of fiber tracks in the distal tendon: the fiber CSAs are smaller, the fiber courses are more curved and in the fibrocartilages, more transversal connections between fibers can be seen. This leads to a bias in the measurements from the fiber tracks. The dimensional measurements might underestimate the *in vivo* dimensions because the applied protocols can bring about tissue shrinkage [4]. The absolute diameters are in the range reported for tendon fibers [16,40,41].

The resulting fiber numbers agree with the dimensional measurements like fiber diameters, nearest neighbor distances, fiber CSAs and tendon CSA. Consequently, the sum of the fiber CSAs decreases towards the insertion due to a decrease in fiber size, but not in fiber number. The larger gaps between the fibers correspond to the presence of rounded cells in this region [42]. In the enthesis rows of chondrones reside between the fibers [14]. Our findings make clear, that the enthesis is not reinforced in terms of a higher number of fibers. However, our measurements of CSA ignore two factors that have to be ruled out, before we can establish a decrease in effective CSA. (1) the fibers in the distal tendon could just be compressed against the *Tuber calcanei* so that the amount of fibrous material is constant along the tendon. In that case, similar numbers of fibrils would be compressed to smaller fiber CSAs in the distal tendon. (2) Apparent fiber CSAs in the proximal tendon could also appear larger due to a higher fiber crimp. To exclude such a bias in the measurements, the distribution of fiber crimp along the tendon has to be quantified.

The calculated tensile stresses are in the range reported for tensile strength of collagen fibers [43,44]. The difference between a proximal and a distal cross-section results from the decrease in fiber CSAs towards the insertion. Higher stress would make tendon ruptures near the insertion more probable if the insertion is not reinforced in other ways. If a decrease in effective CSA was confirmed, other mechanisms that could reinforce the distal tendon, would have to be considered, for example higher strength of the fiber material or recruitment of components between the tendon fibers.

### 4.2 Homogenization of stress by tendon twist

Like in some tendons in rats [15] a subdivision into fascicles was not found in the examined part of the Achilles tendon in mice. Accordingly, branching clusters can be described as the level of organization between the subtendons and the fibers. The lack of endotenon might be a miniaturization effect.

Most of the identified branching clusters comprise a few parallel fibers with low levels of divergence. Such a branching pattern allows sliding between clusters and does not homogenize stress across the tendon. The branching analysis is likely to underestimate the degree of branching because only a part of the fibers in the distal tendon was identified. Nevertheless, our observations do not suggest that the fibers cluster together across the tendon. Barfred [6] observed that the Achilles tendon shows “less interdigitation” in rats than in humans and that the subtendons from the different muscle bellies could be easily separated down to the insertion. We found a similar state in the Achilles tendon in mice.

In mechanical tests on slices of porcine Achilles tendon entheses, Rossetti et al. [16] found that different shares among the fibers at the enthesis are loaded depending on the angle of the force application. They hypothesized that “inter-fiber coupling at the microscale” could contribute to an angle-dependent redistribution of forces. The Achilles tendon in mice seems to lack structures that could lead to the recruitment of different sets of fibers with different joint angles. However, the fiber topology in the fibrocartilages requires further examination with tracking tools that are better adapted to high rates of branching.

Riggin et al. [45] measured fiber orientation with high-frequency ultrasound over the Achilles tendon length of mice and found an angular deviation of 5.6° to 5.8° depending on the method used. The median angular deviations, we measured are higher. This might be due to the fact that our dataset only includes the distal part of the tendon. The increase in angular deviation towards the insertion is attributed to the flattening of the tendon where it curves around the *Tuber calcanei*. Barfred [6] observed that “interdigitation” and tendon twist rarely occur together and hypothesized that they serve alternative functions. We found a high twist and a low level of branching. The function of twist and branching could be clarified by quantification in animals of different size and locomotion type. An obvious mechanical consequence of twisted fiber courses is a homogenization of stress among the fibers because the taut fibers get aligned and thereby displace the slack fibers which in turn experience more stress. The higher orientation of the central fibers of the tendon might lead to subsequent recruitment of the fibers from the center to the margins and demands for a more thorough mechanical analysis – with potential biomimetic implications for rope design.

### 4.3 Curved fibers in the enthesial fibrocartilage

Fibrocartilages are supposed to develop as a response of tendon tissue to compression [15]. The sesamoid fibrocartilage is maintained because of the frequent compression of the tendon against the *Tuber calcanei*. In the enthesial cartilage the local curvature determines how much pressure the fibers exert on the surrounding matrix. In accordance with this idea, we found the highest curvatures in the unmineralized enthesial fibrocartilage and quantified the resulting contact pressure. The presence of pressure-resistant chondrons is required to maintain the curvature. This mutual conditionality between fiber curvature and chondrons might have implications for the regeneration of the fibrocartilage after injury.

The region where tensile stress is expected to peak, overlaps with the region with the highest fiber curvatures. Therefore, equivalent stress is expected to reach a maximum near the insertion. This consideration lends significance to the question whether the insertion is reinforced in terms of a higher strength of the collagen fibers or by other components besides them.

### 4.4 A distinct portion of the tendon

The fibers of the protrusion form a distinct portion of the tendon. By the anterolateral position of this portion within the proximal cross-section, it could correspond to the subtendon formed by the union of the *M. gastrocnemius medialis* and a small part of the *M. gastrocnemius lateralis*. If this is the case, the Achilles tendon rotation is much higher than in humans, where the fibers of the *M. gastrocnemius medialis* insert on the inferior to the lateral side of the insertion area [46] and in rats where the subtendon of the *M. gastrocnemius medialis* inserts laterally and displacements and strain can differ between the subtendons [47].

The fiber tracks do not follow low period undulations of the fibers like fiber crimp. Tortuosity, therefore, reflects the length reserves due to the gross fiber course. Assuming that the length reserves due to low period undulations are equally distributed within the tendon, the fibers of the protrusion are tauter than the rest of the tendon. These findings support the hypothesis that the protrusion might experience a different loading regime, which could explain its formation and shape.

### 4.5 Conclusion

Fiber tracks and transversal sections were extracted from µCTs of the Achilles tendon enthesis in mice. By the analysis of the fiber tracks and the sections, we showed that the enthesis is not reinforced structurally: the fiber number is about constant along the tendon, while the fiber cross-sections become smaller towards the insertion. Correspondingly, tensile stress is expected to reach a maximum near the insertion. At the same time, the tendon fibers are highly curved in the unmineralized entheseal fibrocartilage. The contact pressure exerted onto neighboring chondrons when the tendon transfers tensile stress was estimated. The combination of compressive and tensile stress could render the tendon insertion a weak spot. However, tendon twist was identified as a parameter that could lead to a partial homogenization of stress across the tendon. The branching rate is too low for significant contributions to stress homogenization. Fibers only cluster together in small parallel groups that can slide relative to one another. Fibers inserting at the protrusion of the insertion area were found to form a distinct portion of the tendon and to be tauter than the other fibers. Therefore, the protrusion could experience a different stress regime than the other parts of the insertion area.

### 4.6 Perspectives

The fiber tracks can be used to understand the mechanobiology of fiber formation and for multi-scale simulations of enthesis mechanics. The techniques used for imaging and tracking were proven useful for the analysis of connective tissues. The same approach could be applied to complete muscle-tendon units so that fibers could be tracked from the endomysium of specific muscle motor units to positions at the insertion area.

## Acknowledgments

We hereby thank Markus Löffler for scanning the specimens and introducing us to propagation-based phase-contrast, Jan Giesebrecht and Julien Roussel for testing the feasibility of the fiber tracking approach and giving access to a demo version of the software, Hartmut Witte, Cornelius Schilling, Sebastian Köhring, Jörg Bossert, Martin S. Fischer and the members of the Institute of Zoology and Evolutionary Research for the discussions on the mechanical interpretation of the fiber tracks, Elisabeth Meier for keeping the animals, Katja Felbel and Ingrid Weiß for the laboratory work on the specimens, Jörg U. Hammel, Hans Pohl and Karolin Engelkes for advice with regard to data processing and Dorothea Seeger for proofreading.

## Funding

JS did not receive any specific grant from funding agencies in the public, commercial, or not-for-profit sectors. HS was supported by the Center of Interdisciplinary Prevention of Diseases related to Professional Activities (KIP) funded by the Friedrich-Schiller-University Jena and the ‘Berufsgenossenschaft Nahrungsmittel und Gastgewerbe Erfurt (Germany)’ (BGN).

## Supplement

Supplementary figures can be found online.

[Figure S1]

[Figure S2]

