## Supplemental figures S1, S2 for "Tracking tendon fibers to their insertion – a 3D analysis of the Achilles tendon enthesis in mice"

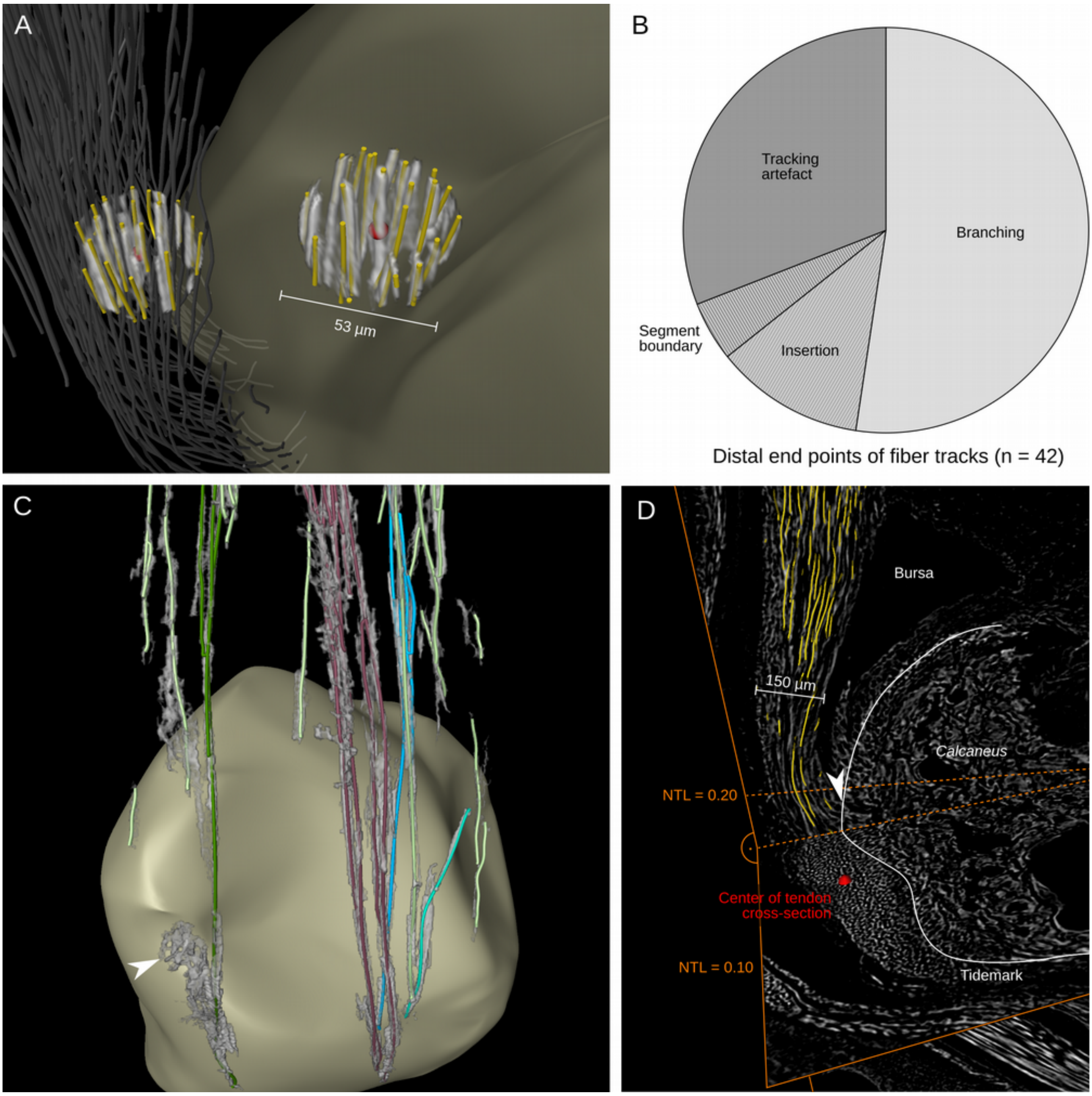


Fig. S1: Evaluation of fiber tracks, endpoints, and branching. (A) Spheres of the volume data and the fiber tracks were extracted around endpoints of fiber tracks in order to evaluate them. (B) Accordingly, a sample of distal end-points of fiber tracks (n = 42) was classified. The majority of these endpoints corresponded to branching points. (C) A rendering of clusters of fiber tracks with the corresponding segments of the volume data shows that distally larger regions are attributed to a cluster (arrowhead). (D) A rendering of fiber tracks together with a sagittal and a transversal slice shows that the proximal-most position where fibers insert (arrowhead) corresponds to a NTL of 0.18.

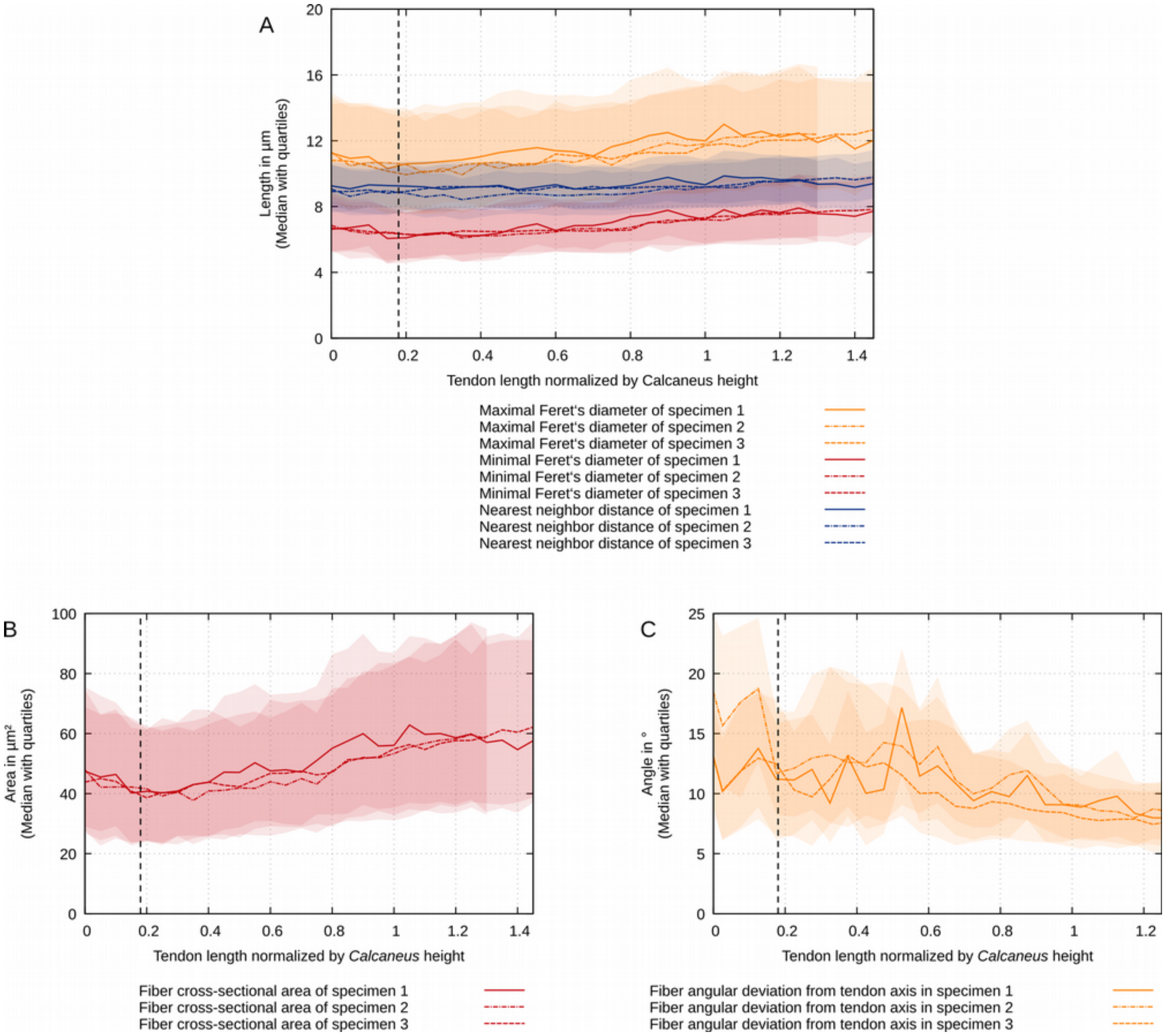


Fig. S2: Fiber sizes and orientations over all three specimens. (A) The medians of the maximal and the minimal Feret's diameter of the fibers in all three specimens decrease towards the insertion. The median nearest neighbor distance of the centroids of the fiber masks stays about constant over NTL. (B) The median CSA of the fibers decreases towards the insertion. (C) The median angular deviation of fiber vectors from the mid-tendon line is is higher near the insertion.
